# Quantifying noise modulation from coupling of stochastic expression to cellular growth: An analytical approach

**DOI:** 10.1101/2022.10.03.510723

**Authors:** Iryna Zabaikina, Zhanhao Zhang, César Nieto, Pavol Bokes, Abhyudai Singh

## Abstract

The overexpression of many proteins can often have a detrimental impact on cellular growth. This expression-growth coupling leads to positive feedback - any increase of intracellular protein concentration reduces the growth rate of cell size expansion that in turn enhances the concentration via reduced dilution. We investigate how such feedback amplifies intrinsic stochasticity in gene expression to drive a skewed distribution of the protein concentration. Our results provide an exact solution to this distribution by analytically solving the Chapman-Kolmogorov equation, and we use it to quantify the enhancement of noise/skewness as a function of expression-growth coupling. This analysis has important implications for the expression of stress factors, where high levels provide protection from stress, but come at the cost of reduced cellular proliferation. Finally, we connect these analytical results to the case of an actively degraded gene product, where the degradation machinery is working close to saturation.

## I. INTRODUCTION

Single cell studies in the past decade have characterized several mechanisms that drive stochasticity in intracellular gene product levels [1]–[7]. These random fluctuations fundamentally affect various biological processes including apoptosis [8], phenotype switching [9], cell lysis by viral infections [10]–[12], and cell differentiation [13]–[17]. As in engineering systems, cells employ feedback and feedforward regulation to modulate stochasticity [18]–[29], attenuating it when fluctuations are deleterious and amplifying it when cell-to-cell heterogeneity is beneficial. The latter case is especially relevant for drug tolerance in microbial/cancer cells, where high intercellular heterogeneity in specific protein levels within an otherwise genetically-identical population allows outlier cells to escape drug treatment [30]–[37].

An important source of expression stochasticity is variability in cellular growth rate [37], [38], and several studies have quantified the interplay between growth rate and gene expression [39]–[44]. For example, the transcription rate of some proteins can increase with growth rate [45]. Here, we will study another perspective on this connection: overexpression of a protein can lead to growth inhibition, as has been shown for several stress factors, enzymes [46], [47], and in the design of synthetic genetic circuits where expression places a significant burden on host cell resources [48]–[50]. The decay in the concentration of a stable protein (i.e., no active degradation) is dominated by dilution from cell growth. Expression-mediated growth inhibition creates positive feedback - any random increase in concentration reduces dilution, which in turn, acts to further increase in concentration [51]–[53].

In this context, we systematically study this expression-growth coupling via a stochastic model where proteins are synthesized in stochastic bursts [2], [54]. In between successive bursts, the protein concentration is diluted continuously as per the nonlinear differential equation.

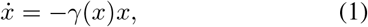

where the dilution rate, in convenient units,

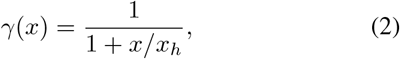

decreases with increasing protein concentration *x*. Here *x*_*h*_ is a positive constant and we refer to 1*/x*_*h*_ as the *feedback strength*. The continuous decay in protein concentration interspersed with discrete burst events constitutes a hybrid system. This hybrid framework has previously provided rich insights on how diverse processes impact the stochastic dynamics of gene product levels [55]–[57]. The key result presented here (Section II) is the exact derivation of the steady-state protein concentration distribution by solving the underlying Chapman-Kolmogorov equation. In the absence of feedback (*x*_*h*_→ ∞ or zero feedback strength), a constant dilution rate leads to a Gamma-distributed protein level [58], [59]. Section III shows, using the moment analysis perspective, how the feedback in growth rate affects the protein statistics for an arbitrary intrinsic noise.

An analogous problem arises in the context of an unstable protein (i.e., a protein that is actively degraded), but where the degradation machinery/enzymes operate close to saturation. To study this problem, we formulate a similar stochastic model, but here the protein level is modeled as an integer-valued random process and degradation is a discrete event that occurs at a probabilistic rate *γ*(*x*)*x*. In Section IV, taking a moment analysis approach, we show how steady-state moments of protein population counts can be obtained exactly despite the fact that the rates are rational functions of *x*. The procedure used to obtain exact solutions of moments can also be used for other stochastic dynamical systems, where saturation effects are modeled through rational functions.

## II. MODELING GROWTH-INHIBITION MEDIATED FEEDBACK (CONTINUOUS PROTEIN LEVEL)

Consider a stable protein with concentration *x*(*t*) ∈ℝ^+^; at time *t* diluted as per (1)-(2). Protein synthesis events occur randomly at a rate *k*_*x*_ and each event creates a “jump” or burst in the concentration

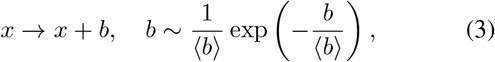

where the burst size *b* is an i.i.d. random variable that follows an exponential distribution with mean⟨*b*⟩. Here and throughout the paper we use ⟨⟩ to denote the expected value of random variables and random processes. It is important to point out that for the model to be physiologically relevant, the parameters should satisfy

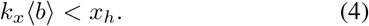

To see this note that *k*_*x*_ ⟨*b*⟩ is the average protein synthesis rate, while the total dilution rate

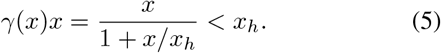

Thus, *k*_*x*_ ⟨*b*⟩ has to be less than the maximum net decay rate *x*_*h*_, else the concentration will increase unboundedly over time. The overall model schematic is shown in Fig. 1A. Fig. 1C. shows stochastic trajectories of *x*. Fig. 1D. presents dilution rates *γ*(*x*) for different values *x*_*h*_ under the same burst events (Fig. 1B).

**Fig. 1.**
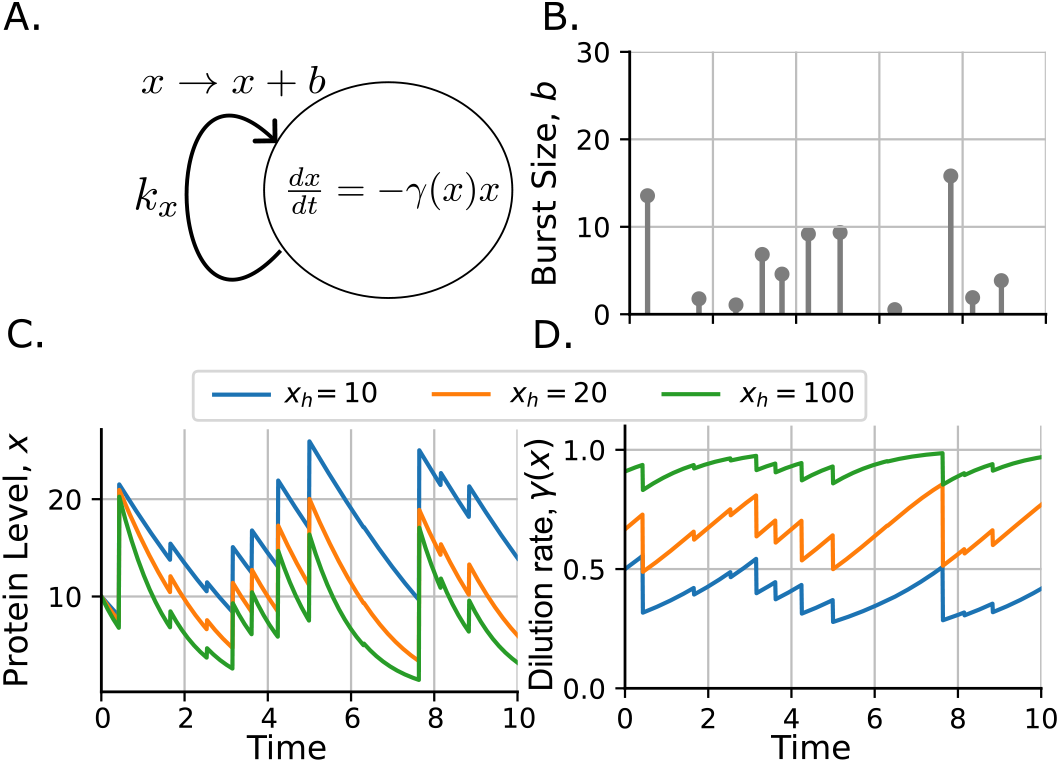
Illustration of feedback model coupling continuous gene expression with dilution. A. Diagram of the stochastic hybrid system that describes protein dynamics. Protein synthesis *x* ∈ ℝ^+^ is a jump process, that increases by a random burst size *b* ∈ ℝ^+^ each time a burst event occurs. Between consecutive burst events, the protein dilutes following the differential equation inside the ellipse. B. Random size protein burst events that occur as per a Poisson rate. C. Protein concentration dynamics considering different halfsaturation concentrations *x*_*h*_ D. Dynamics of the dilution rate as given by (2) for the protein levels shown in C.

### A. Derivation of the steady-state distribution

Let *p*(*x, t*) denote the probability density function (PDF) of protein concentration *x* at time *t*. Based on the model formulation in Fig. 1, the time evolution of *p*(*x, t*) is described by the Chapman-Kolmogorov equation [58], [60],

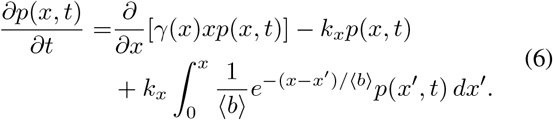

The steady-state protein PDF is given by (See Appendix A for detail),

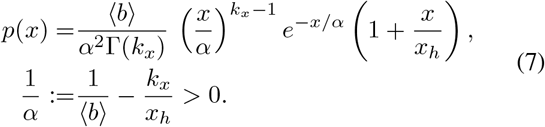

In the absence of feedback (i.e, *x*_*h*_→ ∞ that corresponds to a constant dilution rate *γ*(*x*) = 1), *p*(*x*) reduces to a Gamma distribution

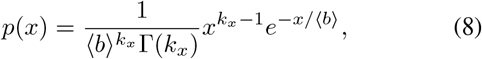

as has been previously derived and also found consistent with experimentally observed distributions [58], [59].

Multiplying (7) by *x, x*^2^ or *x*^3^ correspondingly and integration it on the interval [0, ∞) give the first three orders steady state moment,

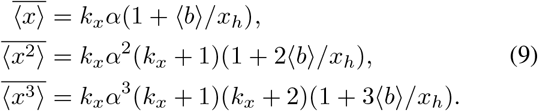

Fig. 2 presents typical stochastic trajectories of protein concentration as well as a comparison of PDF between the simulated (histogram) and analytical (solid lines) results. Analytical results for constant dilution rate (Fig. 2A) and concentration-dependent feedback dilution rate (Fig. 2B) show good match with stochastic simulation results. Fig. 3 presents a comparison of PDFs between different feedback strengths (1*/x*_*h*_). As we increase the feedback strength, the tendency of skew in the distribution is enhanced.

**Fig. 2.**
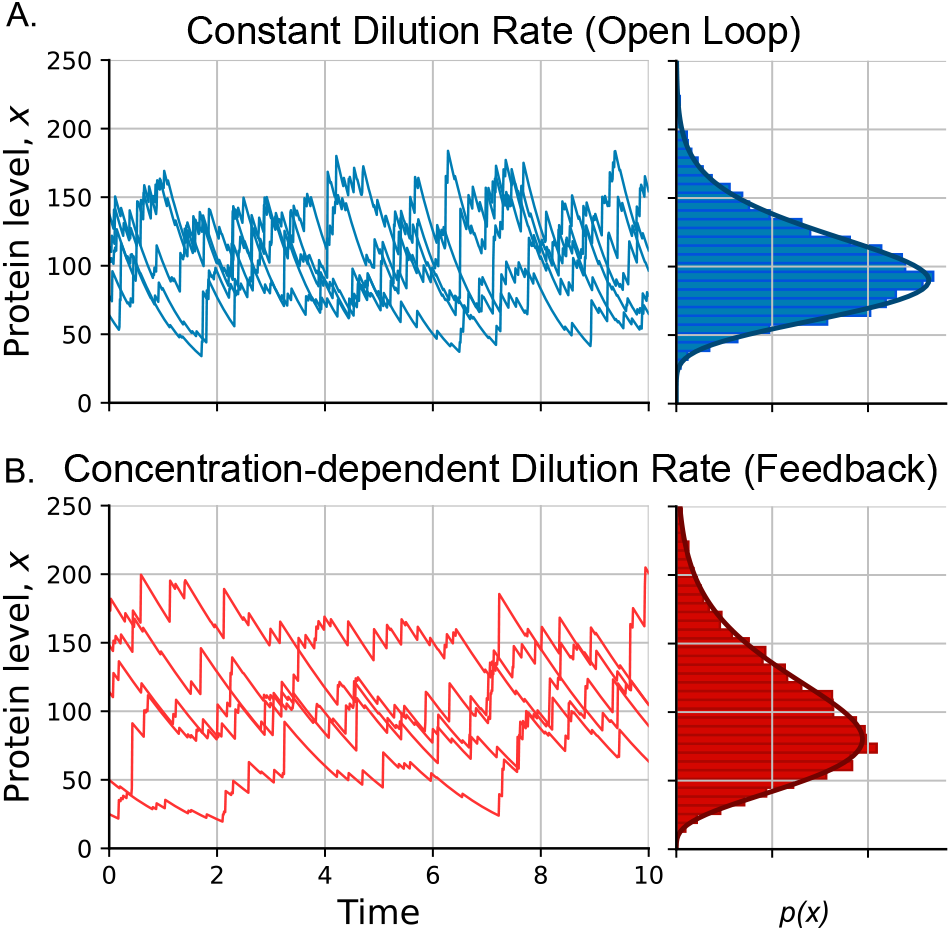
Protein trajectories (left) and steady-state distributions (right). A. Constant dilution rate. B. Concentration-dependent dilution rate. The PDF drawn with solid line corresponds to the analytical expression, while the histogram is the result of stochastic sim<u>ula</u>tions. (Parameters: ⟨*b*⟩ = 10, *x*_*h*_ = 100, *k*_*x*_ is calculated such as ⟨*x*⟩ = 100 in both cases).

**Fig. 3.**
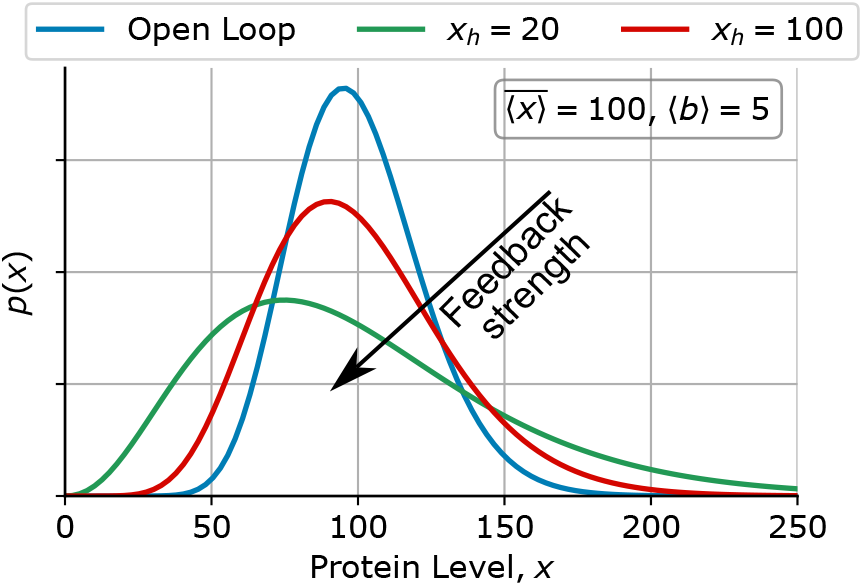
Comparison of steady-state distributions. The protein concentration probability distribution function (7) is plotted with different feedback strengths (1*/x*_*h*_). Other parameters taken as ⟨*b*⟩ = 5, *k*_*x*_ is calculated so as to fix the steady-state mean level 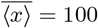).

## III. MOMENT ANALYSIS METHOD FOR CONTINUOUS PROTEIN LEVEL

In the above section, we derived the protein concentration PDF for the case where the protein bursts *b* follow an exponential distribution. In the general case, we measure the moments of *b* with no particular assumption of its distribution. We next show how a moment dynamics approach can be employed to derive the steady-state statistical moments of *x*(*t*) without assumptions about the PDF of *b*.

### A. Constant dilution rate

To introduce the moment analysis method, let us consider that the burst size follows any arbitrary PDF *ν*(*b*). If during each burst, *x* evolves as (1), the time derivative of the expected value of any arbitrary function *φ*(*x*) satisfies [61]

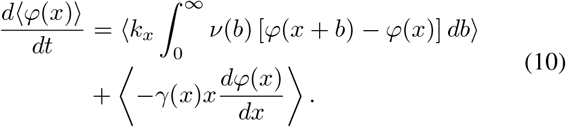

Considering *γ*(*x*) = 1 for the no-feedback case and setting *φ*(*x*) = *x, x*^2^, and *x*^3^ in (10) yields the following system of differential equations

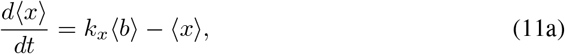

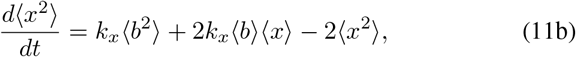

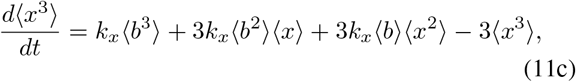

Where ⟨*b*^*n*^⟩ =∫ *b*^*n*^*ν*(*b*)*db*.

Setting the left-hand-side of (11) to zero we obtain the steady-state moments 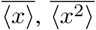, and 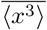 as

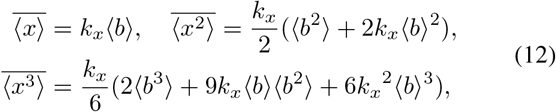

respectively. Using these uncentered moments one can quantify the noise in protein concentration via the steady-state Fano factor (variance divided by mean)

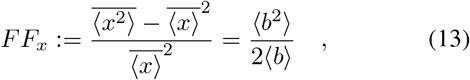

and the skewness

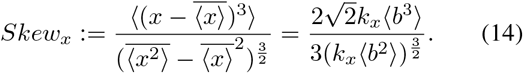

When *b* is exponentially distributed, *FF*_*x*_ and *Skew*_*x*_ reduce to

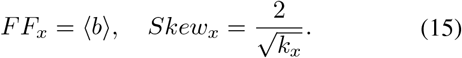

### B. Concentration-dependent dilution rate

Now, let us consider the scenario where the dilution rate *γ*(*x*) depends on *x* as in (2). Using (10), we obtain the equations describing the moment dynamics

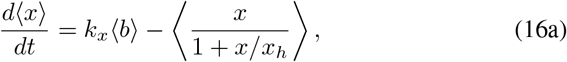

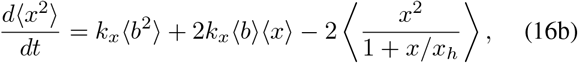

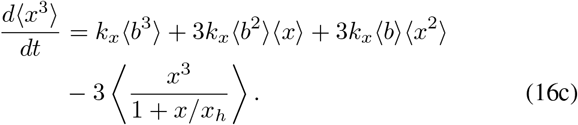

Given the nonlinearity arising via the rational decay rate, (16a) by itself cannot be solved to obtain a formula for 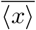. However, it turns out that this mean is obtained by simultaneously solving (16a)-(16b). To see this, we note from (16a)

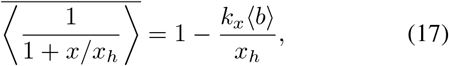

where the 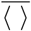 symbol denotes the expected value at steadystate. Using the fact that

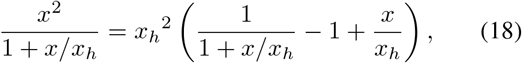

we obtain from (16b) at steady-state

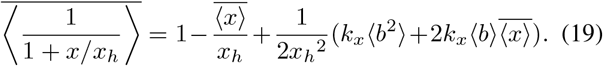

By setting (17) equal to (19), one can solve for the steadystate mean level

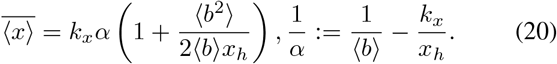

As expected, in the limit *x*_*h*_→ ∞, *α*→⟨*b*⟩ this mean converges to the no-feedback case (12).

Using a similar approach one can use the first three moment equations (16) to obtain 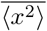. To see this, by writing

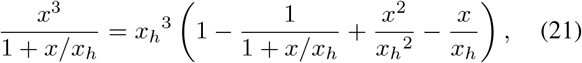

we obtain from (16c),

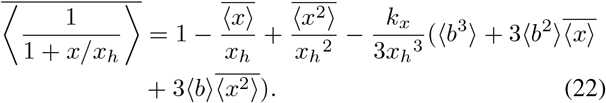

And setting (19) equal to (22) results in

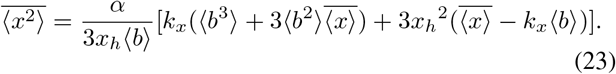

Following along this direction, to derive the third order steady-state moment one requires the fourth order moment dynamics,

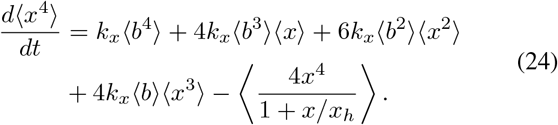

By performing a fraction decomposition of the last term in (24) and expressing it in terms of 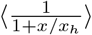, and substituting its steady-state value from (22) back in (24) leads to

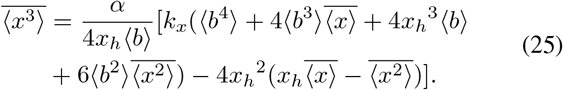

These steady-state moments can be used to obtain the Fano factor and skewness,

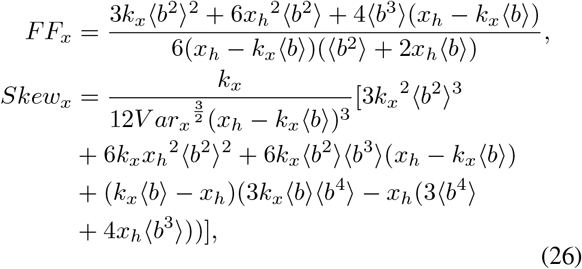

respectively, where *V ar*_*x*_ is the variance of steady-state protein concentration,

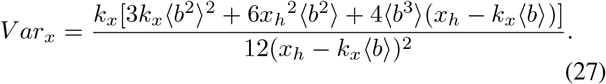

For the case where the burst size follows an exponential distribution,

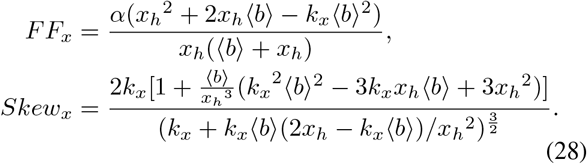

In the limit of no feedback (*x*_*h*_→ ∞, *α*→⟨*b*⟩), (28) converges to (15).To analyze the results, in Fig. 4A, we present how the noise in protein level changes with burst size ⟨*b*⟩. For this plot, we tune *k*_*x*_ while keeping 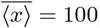 and *x*_*h*_ = 100. In the absence of feedback, *FF*_*x*_ = ⟨*b*⟩. In the presence of feedback, the noise in protein concentration increases with burst size ⟨*b*⟩. Fig. 4B shows how the noise in protein is amplified by increasing feedback strength (1*/x*_*h*_). A similar analysis is made with skewness in Fig. 4C and D.

**Fig. 4.**
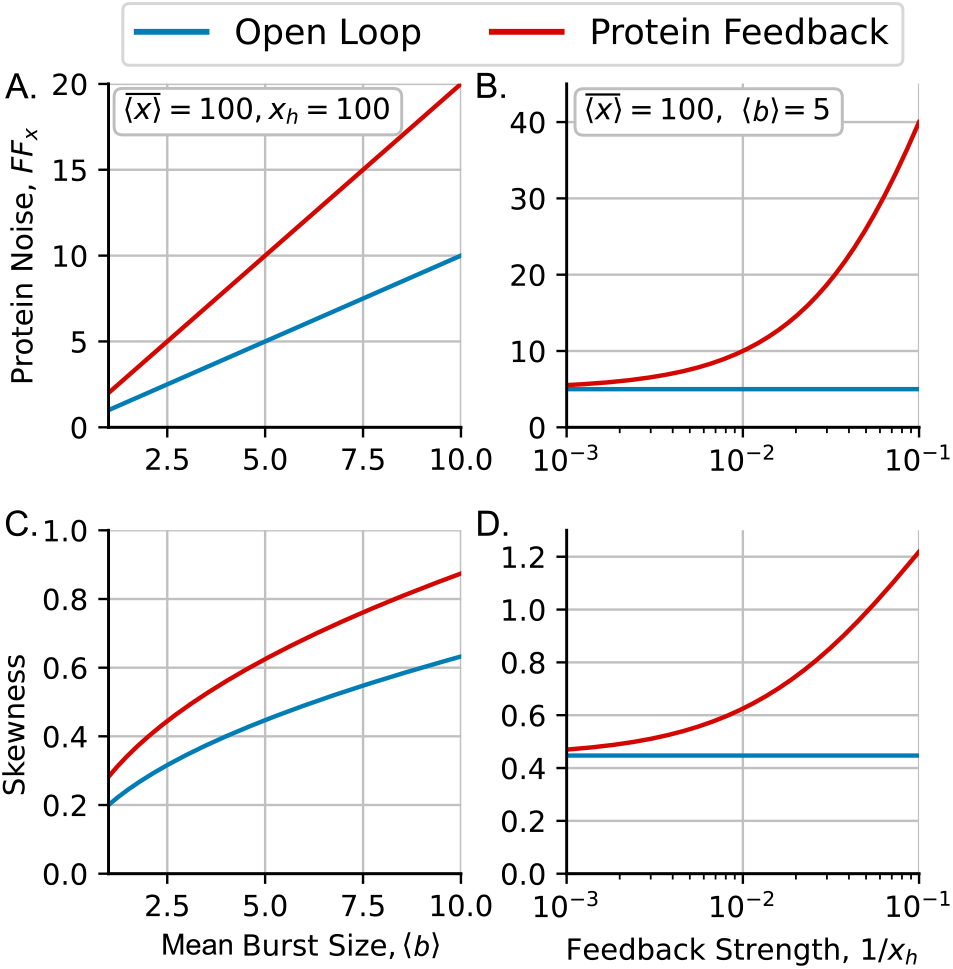
Amplification in protein noise and skewness by the coupling of expression level to cellular growth. A. Protein noise plotted via the steady-state Fano factor *FF*_*x*_ against the mean burst size ⟨*b*⟩. B. Protein noise *FF*_*x*_ vs. the feedback strength (1*/x*_*h*_). C. Protein Skewness vs the mean burst size. D. skewness in protein concentration vs feedback strength (1*/x*_*h*_). All figure we assume an exponential distribution for *b*. (Parameters: 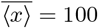, *x*_*h*_ = 100 in A and C, and ⟨*b*⟩ = 5 in B and D.)

## IV. MOMENTS ANALYSIS FOR SATURATION IN DEGRADATION MACHINERY (DISCRETE PROTEIN LEVEL)

In this section, we explore an alternative scenario where the protein is actively degraded by an enzyme that is present in limiting amounts, resulting in a degradation rate given by

(1). To capture the stochastic dynamics we now consider an analogous integer-valued random process *x*(*t*) with two key differences (Fig.5):

**Fig. 5.**
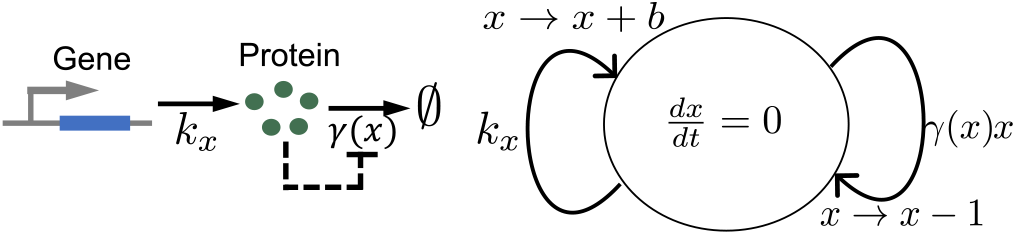
Diagram of the stochastic system that describes protein synthesis and active degradation. Feedback arising from degradation rate *γ*(*x*) decreases with increasing protein level. Protein synthesis *x* ∈ ℕ is modeled as a jump process which increases by a random burst size *b* ∈ ℕ each time a burst event occurs. Degradation events occur with rate *γ*(*x*)*x* and reduce *x* by one.

1) Upon a burst event, *x*(*t*) increases by *b* where the burst size now follows an arbitrary integer-valued distribution 𝕡(*b*). A standard distribution often used for burst size in literature is a geometric distribution [62].

2) The degradation is modeled as a probabilistic event that occurs with rate *xγ*(*x*), and protein numbers decrease by one when the event occurs.

For this discrete system, the time evolution of moments can be obtained using the generator [63]

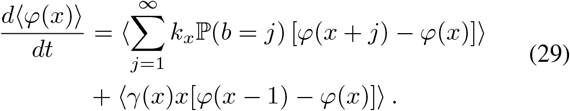

### A. Constant degradation rate

The moment dynamics for the no-feedback case (*γ*(*x*) =

1) are

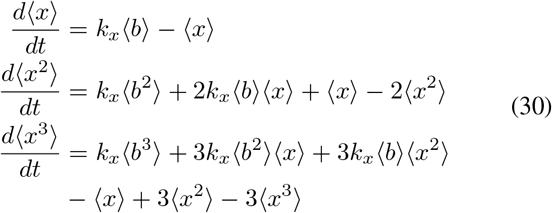

which yields the following steady-state moments, Fano factor, and skewness,

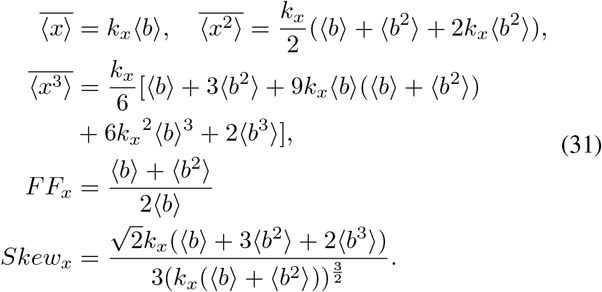

When the burst size is geometric-distributed, ℙ (b) = (1 −f)bf with f = 1/(1 + ⟨b⟩) we get the Fano factor and skewness

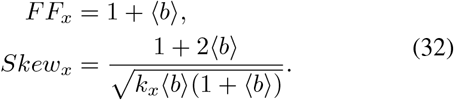

and a straightforward comparison confirms a higher *FF*_*x*_ in the discrete formulation compared to its continuous counterpart in (15). See the dashed line in Fig. 6A.

**Fig. 6.**
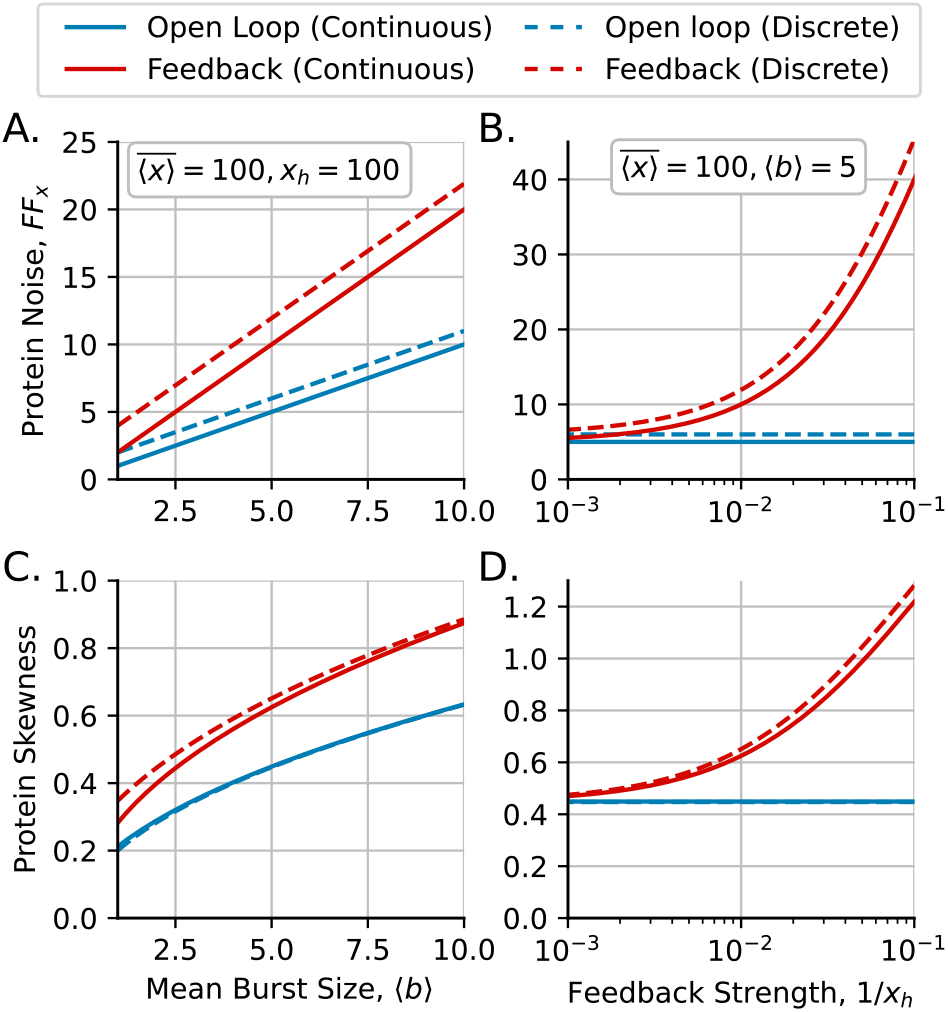
Amplification in protein noise and skewness due to saturation in degradation machinery. A. Protein noise *FF*_*x*_ vs mean burst size ⟨*b*⟩ with the same mean protein level. B. Protein noise *FF*_*x*_ as the feedback strength (1*/x*_*h*_) increases. C. Skewness of the protein concentration vs mean burst size ⟨*b*⟩. D. Skewness vs feedback strength (1*/x*_*h*_). We assume geometric distribution for *b*. Statistics for discrete protein (dashed lines) are compared to the statistics considering continuous protein levels (continuous lines). Results with the open loop protein synthesis (blue) are also compared to the result considering feedback (r<u>ed).</u> (Parameters: *x*_*h*_ = 100 in A and C, ⟨*b*⟩ = 5 in B and D, 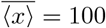, and *k*_*x*_ is calculated such as 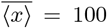 in both cases)

### B. Protein feedback in degradation rate

For feedback degradation, *γ*(*x*) = 1*/*(1 + *x/x*_*h*_), we can obtain steady-state moments by performing a similar method in (17)-(23) (See Appendix B for detail). The Fano factor and the skewness become,

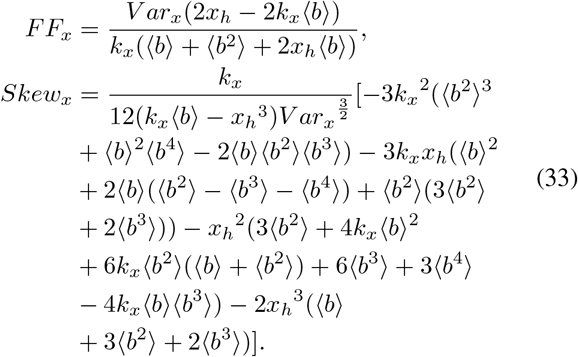

where,

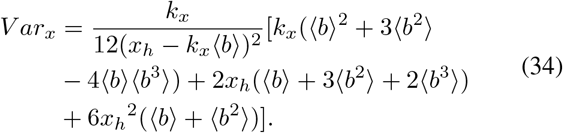

For the case where *b* follows a geometric distribution,

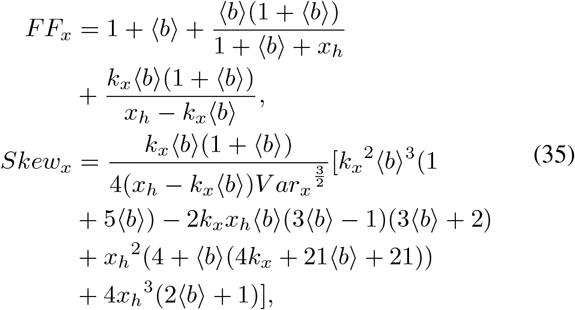

where,

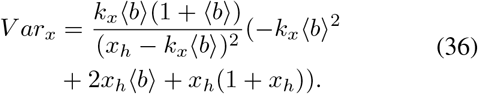

In the limit of absence of feedback (*x*_*h*_→ ∞, *γ*(*x*) → 1), *FF*_*x*_ and *Skew*_*x*_ in (35) become (32).

Fig. 6 presents, in a similar way as Fig. 4, the Fano factor and skewness for continuous dilution (solid line) and discrete degradation (dashed line). In Fig. 6A and B, the noise in protein level (Fano factor) is plotted against the burst size (⟨*b*⟩) and feedback strength (1*/x*_*h*_), respectively. Both cases have qualitatively similar trends, but the discrete case has a larger Fano factor regardless of the presence of feedback. Fig. 6C and D show the comparison of skewness. In the absence of feedback, the skewness of both cases has similar values. However, in the presence of feedback, the skewness of the discrete case is larger than that of the continuous case.

## V. CONCLUSION

In this contribution, we have systematically explored how the expression-growth coupling implements a positive feedback in gene expression (Fig. 1). We studied the impact of this feedback on the amplification of noise at the level of a given protein. Our exact derivation of the steadystate protein level distribution (Fig. 2) quantifies both the enhancement of noise and the skewness as a function of the feedback strength (Fig. 4). Having this analytically predicted distribution is useful for inferring parameters by fitting it to data on single-cell stress factor levels that are known to inhibit cellular proliferation [49], [50]. Our results show that such forms of feedback could lead to a larger fraction of outlier cells with high expression levels (Fig. 3). This stochastically primed outlier cell subpopulation has lower fitness in terms of proliferation, but can survive detrimental changes in extracellular environments.

In the context of cancer, the survival of a rare subpopulation of stochastically prime drug-tolerant cells in response to targeted therapy often drives the development of drug resistance [64]. We further extended these results with a discrete-state formulation of protein synthesis where the degradation rate is inversely related to its expression level.

This work paves the way for several directions of future investigations. For example, given a certain frequency of drug treatment regimes that favor survival of outlier cells, is there an optimal feedback strength that maximizes the populations long-term fitness in fluctuating environments? The analysis here is based on following a single cell over time and this naturally leads to exploring the protein distribution in an expanding cell population using a combination of simulations and population balance equations [65]–[67].

APPENDIX

### A. Derivation of steady-state distribution

Let *p*(*x, t*) denote the probability density function of protein concentration at time *t*. Based on the model formulation in Fig. 1, the time evolution of *p*(*x, t*) is described by the Chapman-Kolmogorov equation,

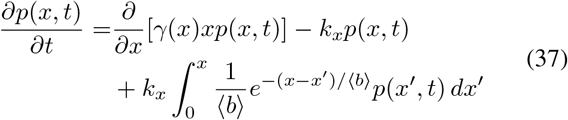

[58], [60]. In order to find the stationary distribution, we rewrite it in the form of a partial integro-differential equation,

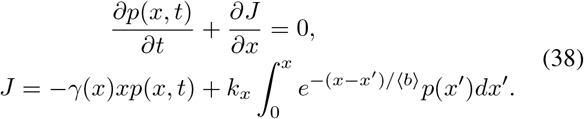

At steady state, *∂p/∂t* = 0, this yields the Volterra integral equation[60],

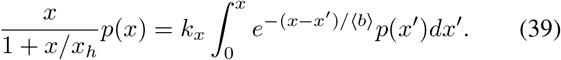

We present the method of solving (39) to obtain the steadystate protein concentration distribution based on the Laplace transform. In case we want to use Laplace transform approach to solve equation (39), we perform a rearrangement in the following way,

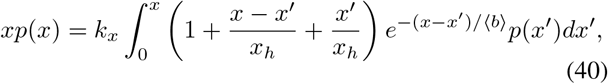

after which the right side of equation becomes the sum of three distinct convolutions. Define an image *P* (*s*) as a function of a Laplace variable *s*,

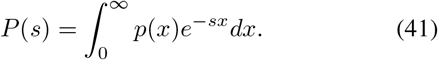

Applying the Laplace transform to (40), we find

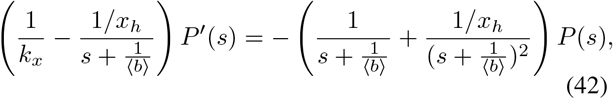

which becomes a separable differential equation

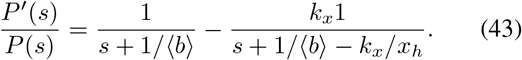

The general solution of (43) is given by

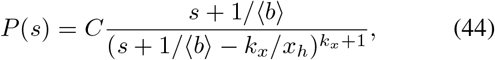

where *C* is an arbitrary constant. Note that the right hand side is a power function of the Laplace variable *s*, which is shifted by value

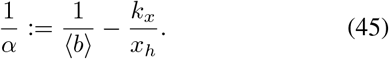

In order to return to the original function *p*(*x*), we take inverse Laplace transform applied to the general solution by using the following relation,

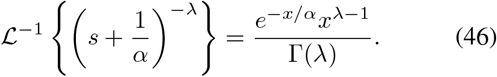

We obtain the general solution of (44),

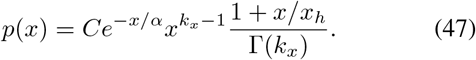

We set *C* so that the integral of the right side is equal to one (i.e., *p*(*x*) is probability density function on the non-negative domain), the steady state probability density function of protein concentration is given by,

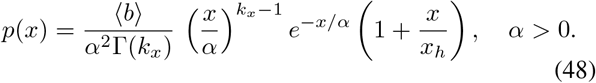

### B. Moment analysis of feedback active degradation

For feedback degradation *γ*(*x*) = *x/*(1 + *x/x*_*h*_), The first three order moment can be obtained by setting *φ*(*x*) = *x, x*^2^, and *x*^3^ in (29), respectively,

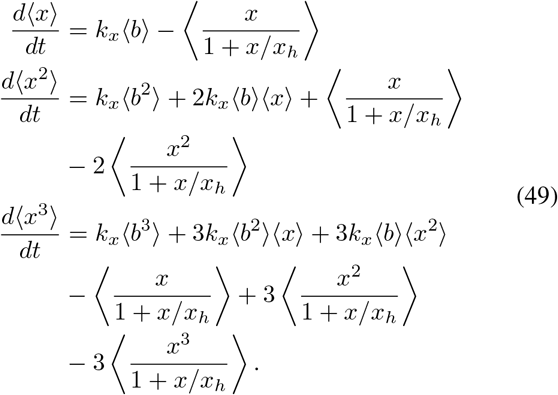

By performing similar approach in (17)-(23), we can obtain the steady-state moment,

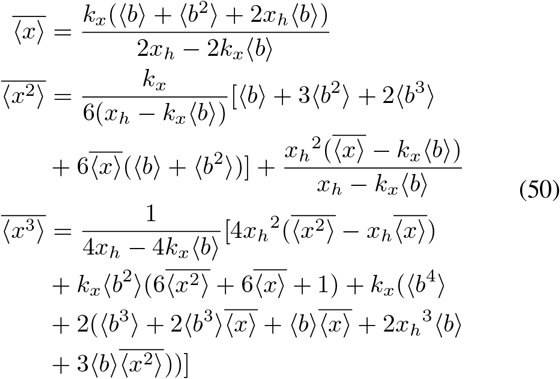

The Fano factor and the skewness become,

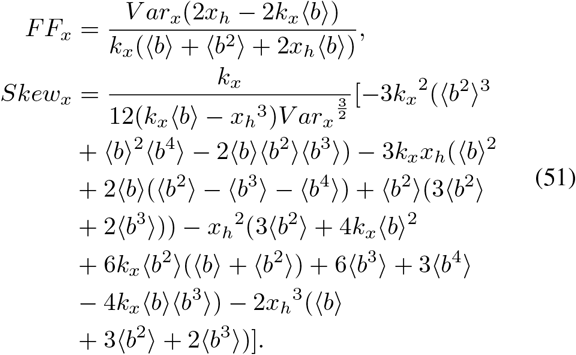

where,

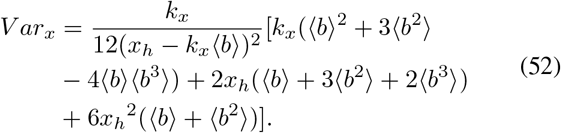

## REFERENCES

[1] M. B. Elowitz, A. J. Levine, E. D. Siggia, and P. S. Swain, “Stochastic gene expression in a single cell,” Science, vol. 297, no. 5584, pp. 1183–1186, 2002.

[2] T. Jia and R. V. Kulkarni, “Intrinsic noise in stochastic models of gene expression with molecular memory and bursting,” Physical review letters, vol. 106, no. 5, p. 058102, 2011.

[3] L. C. Fraser, R. J. Dikdan, S. Dey, A. Singh, and S. Tyagi, “Reduction in gene expression noise by targeted increase in accessibility at gene loci,” Proceedings of the National Academy of Sciences, vol. 118, 2021.

[4] Raj, C. Peskin, D. Tranchina, D. Vargas, and S. Tyagi, “Stochastic mRNA synthesis in mammalian cells,” PLOS Biology, vol. 4, p. e309, 2006.

[5] D. M. Suter, N. Molina, D. Gatfield, K. Schneider, U. Schibler, and F. Naef, “Mammalian genes are transcribed with widely different bursting kinetics,” Science, vol. 332, pp. 472–474, 2011.

[6] B. Schwanhäusser, D. Busse, N. Li, G. Dittmar, J. Schuchhardt, J. Wolf, W. Chen, and M. Selbach, “Global quantification of mammalian gene expression control,” Nature, vol. 473, no. 7347, pp. 337– 342, 2011.

[7] J. Paulsson, “Models of stochastic gene expression,” Physics of life reviews, vol. 2, no. 2, pp. 157–175, 2005.

[8] J. Roux, M. Hafner, S. Bandara, J. J. Sims, H. Hudson, D. Chai, and P. K. Sorger, “Fractional killing arises from cell-to-cell variability in overcoming a caspase activity threshold,” Molecular systems biology, vol. 11, no. 5, p. 803, 2015.

[9] E. M. Ozbudak, M. Thattai, H. N. Lim, B. I. Shraiman, and Van Oudenaarden, “Multistability in the lactose utilization network of escherichia coli,” Nature, vol. 427, no. 6976, pp. 737–740, 2004.

[10] K. Rijal, A. Prasad, A. Singh, and D. Das, “Exact distribution of threshold crossing times for protein concentrations: Implication for biological timekeeping,” Physical Review Letters, vol. 128, no. 4, p. 048101, 2022.

[11] K. R. Ghusinga, J. J. Dennehy, and A. Singh, “First-passage time approach to controlling noise in the timing of intracellular events,” Proceedings of the National Academy of Sciences, vol. 114, pp. 693– 698, 2017.

[12] S. Kannoly, T. Gao, S. Dey, N. Wang, A. Singh, and J. J. Dennehy, “Optimum threshold minimizes noise in timing of intracellular events,” Iscience, vol. 23, p. 101186, 2020.

[13] M. A. Coomer, L. Ham, and M. P. Stumpf, “Noise distorts the epigenetic landscape and shapes cell-fate decisions,” Cell Systems, vol. 13, no. 1, pp. 83–102, 2022.

[14] T. M. A. Neildez-Nguyen, A. Parisot, C. Vignal, P. Rameau, D. Stockholm, J. Picot, V. Allo, C. Le Bec, C. Laplace, and A. Paldi, “Epigenetic gene expression noise and phenotypic diversification of clonal cell populations,” Differentiation, vol. 76, pp. 33–40, 2008.

[15] T. M. Norman, N. D. Lord, J. Paulsson, and R. Losick, “Stochastic switching of cell fate in microbes,” Annual Review of Microbiology, vol. 69, pp. 381–403, 2015.

[16] A. Singh, “Stochastic analysis of genetic feedback circuit controlling HIV cell-fate decision,” Proc. of the 51st IEEE Conf. on Decision and Control, Maui, Hawaii, pp. 4918–4923, 2012.

[17] T. Akanuma, C. Chen, T. Sato, R. M. Merks, and T. N. Sato, “Memory of cell shape biases stochastic fate decision-making despite mitotic rounding,” Nature communications, vol. 7, p. 11963, 2016.

[18] L. S. Weinberger, J. Burnett, J. Toettcher, A. Arkin, and D. Schaffer, “Stochastic gene expression in a lentiviral positive-feedback loop: HIV-1 Tat fluctuations drive phenotypic diversity,” Cell, vol. 122, pp. 169–182, 2005.

[19] S. Hooshangi and R. Weiss, “The effect of negative feedback on noise propagation in transcriptional gene networks,” Chaos: An Interdisciplinary Journal of Nonlinear Science, vol. 16, 2006.

[20] Y. Dublanche, K. Michalodimitrakis, N. Kummerer, M. Foglierini, and L. Serrano, “Noise in transcription negative feedback loops: simulation and experimental analysis,” Molecular Systems Biology, vol. 2, p. 41, 2006.

[21] N. Kumar, T. Platini, and R. V. Kulkarni, “Exact Distributions for Stochastic Gene Expression Models with Bursting and Feedback,” Physical Review Letters, vol. 113, p. 268105, 2014.

[22] A. Singh and J. P. Hespanha, “Optimal feedback strength for noise suppression in autoregulatory gene networks,” Biophysical Journal, vol. 96, pp. 4013–4023, 2009.

[23] A. Singh, “Negative feedback through mRNA provides the best control of gene-expression noise,” IEEE Transactions on Nanobioscience, vol. 10, pp. 194–200, 2011.

[24] Y. Tao, X. Zheng, and Y. Sun, “Effect of feedback regulation on stochastic gene expression,” Journal of Theoretical Biology, vol. 247, pp. 827–836, 2007.

[25] M. Voliotis and C. G. Bowsher, “The magnitude and colour of noise in genetic negative feedback systems,” Nucleic Acids Research, 2012.

[26] T. Kang, T. Quarton, C. M. Nowak, K. Ehrhardt, A. Singh, Y. Li, and L. Bleris, “Robust filtering and noise suppression in intragenic mirna-mediated host regulation,” iScience, vol. 23, no. 10, p. 101595, 2020.

[27] P. Bokes, A. Borri, P. Palumbo, and A. Singh, “Mixture distributions in a stochastic gene expression model with delayed feedback: a wkb approximation approach,” Journal of Mathematical Biology, vol. 81, pp. 343–367, 2020.

[28] C. Tan, P. Marguet, and L. You, “Emergent bistability by a growth-modulating positive feedback circuit,” Nature chemical biology, vol. 5, no. 11, pp. 842–848, 2009.

[29] A. Borri, P. Palumbo, and A. Singh, “Noise propagation in metabolic pathways: the role of growth-mediated feedback,” in 2020 59th IEEE Conference on Decision and Control (CDC). IEEE, 2020, pp. 4610– 4615.

[30] S. M. Shaffer, M. C. Dunagin, S. R. Torborg, E. A. Torre, B. Emert, Krepler, M. Beqiri, K. Sproesser, P. A. Brafford, M. Xiao, E. Eggan, N. Anastopoulos, C. A. Vargas-Garcia, A. Singh, K. L. Nathanson, M. Herlyn, and A. Raj, “Rare cell variability and drug-induced reprogramming as a mode of cancer drug resistance,” Nature, vol. 546, pp. 431–435, 2017.

[31] S. M. Shaffer, B. L. Emert, R. A. R. Hueros, C. Cote, G. Harmange, L. Schaff, A. E. Sizemore, R. Gupte, E. Torre, A. Singh et al., “Memory sequencing reveals heritable single-cell gene expression programs associated with distinct cellular behaviors,” Cell, vol. 182, pp. 947–959, 2020.

[32] I. E. Meouche, Y. Siu, and M. J. Dunlop, “Stochastic expression of a multiple antibiotic resistance activator confers transient resistance in single cells,” Scientific Reports, vol. 6, p. 19538, 2016.

[33] C. A. Chang, J. Jen, S. Jiang, A. Sayad, A. S. Mer, K. R. Brown, M. Nixon, A. Dhabaria, K. H. Tang, D. Venet et al., “Ontogeny and vulnerabilities of drug-tolerant persisters in her2+ breast cancer,” Cancer discovery, 2021.

[34] J. J. Lee, S.-K. Lee, N. Song, T. O. Nathan, B. M. Swarts, S.-Y. Eum, S. Ehrt, S.-N. Cho, and H. Eoh, “Transient drug-tolerance and permanent drug-resistance rely on the trehalose-catalytic shift in mycobacterium tuberculosis,” Nature communications, vol. 10, pp. 1– 12, 2019.

[35] E. A. Libby, S. Reuveni, and J. Dworkin, “Multisite phosphorylation drives phenotypic variation in (p) ppgpp synthetase-dependent antibiotic tolerance,” Nature communications, vol. 10, pp. 1–10, 2019.

[36] I. Levin-Reisman, I. Ronin, O. Gefen, I. Braniss, N. Shoresh, and N. Q. Balaban, “Antibiotic tolerance facilitates the evolution of resistance,” Science, vol. 355, pp. 826–830, 2017.

[37] D. J. Kiviet, P. Nghe, N. Walker, S. Boulineau, V. Sunderlikova, and S. J. Tans, “Stochasticity of metabolism and growth at the single-cell level,” Nature, vol. 514, no. 7522, pp. 376–379, 2014.

[38] P. Thomas, G. Terradot, V. Danos, and A. Y. Weiße, “Sources, propagation and consequences of stochasticity in cellular growth,” Nature communications, vol. 9, no. 1, pp. 1–11, 2018.

[39] M. Scott, C. W. Gunderson, E. M. Mateescu, Z. Zhang, and T. Hwa, “Interdependence of cell growth and gene expression: origins and consequences,” Science, vol. 330, no. 6007, pp. 1099–1102, 2010.

[40] M. Kafri, E. Metzl-Raz, G. Jona, and N. Barkai, “The cost of protein production,” Cell reports, vol. 14, no. 1, pp. 22–31, 2016.

[41] L. H. Krah and R. Hermsen, “The effect of natural selection on the propagation of protein expression noise to bacterial growth,” PLoS computational biology, vol. 17, no. 7, p. e1009208, 2021.

[42] S. Dey and A. Singh, “Stochastic analysis of feedback control by molecular sequestration,” in 2019 American Control Conference (ACC). IEEE, 2019, pp. 4466–4471.

[43] J. Feng, D. A. Kessler, E. Ben-Jacob, and H. Levine, “Growth feedback as a basis for persister bistability,” Proceedings of the National Academy of Sciences, vol. 111, pp. 544–549, 2014.

[44] S. Klumpp and T. Hwa, “Bacterial growth: global effects on gene expression, growth feedback and proteome partition,” Current Opinion in Biotechnology, vol. 28, pp. 96–102, 2014.

[45] M. P. Swaffer, J. Kim, D. Chandler-Brown, M. Langhinrichs, G. K. Marinov, W. J. Greenleaf, A. Kundaje, K. M. Schmoller, and J. M. Skotheim, “Transcriptional and chromatin-based partitioning mechanisms uncouple protein scaling from cell size,” Molecular Cell, vol. 81, no. 23, pp. 4861–4875, 2021.

[46] E. Dekel and U. Alon, “Optimality and evolutionary tuning of the expression level of a protein,” Nature, vol. 436, no. 7050, pp. 588– 592, 2005.

[47] D. Molenaar, R. Van Berlo, D. De Ridder, and B. Teusink, “Shifts in growth strategies reflect tradeoffs in cellular economics,” Molecular systems biology, vol. 5, no. 1, p. 323, 2009.

[48] S. Vadia and P. A. Levin, “Growth rate and cell size: a re-examination of the growth law,” Current opinion in microbiology, vol. 24, pp. 96– 103, 2015.

[49] O. Patange, C. Schwall, M. Jones, C. Villava, D. A. Griffith, Phillips, and J. C. Locke, “Escherichia coli can survive stress by noisy growth modulation,” Nature communications, vol. 9, no. 1, pp. 1–11, 2018.

[50] G. Harmange, R. A. R. Hueros, D. L. Schaff, B. L. Emert, M. M. Saint-Antoine, S. Nellore, M. E. Fane, G. M. Alicea, A. T. Weeraratna, Singh et al., “Disrupting cellular memory to overcome drug resistance,” bioRxiv, 2022.

[51] J. R. Melendez-Alvarez and X.-J. Tian, “Emergence of qualitative states in synthetic circuits driven by ultrasensitive growth feedback,” PLOS Computational Biology, vol. 18, no. 9, p. e1010518, 2022.

[52] H. Y. Kueh, A. Champhekar, S. L. Nutt, M. B. Elowitz, and E. V. Rothenberg, “Positive feedback between pu. 1 and the cell cycle controls myeloid differentiation,” Science, vol. 341, no. 6146, pp. 670– 673, 2013.

[53] C. Jia, P. Xie, M. Chen, and M. Q. Zhang, “Stochastic fluctuations can reveal the feedback signs of gene regulatory networks at the single-molecule level,” Scientific reports, vol. 7, no. 1, pp. 1–9, 2017.

[54] V. Shahrezaei and P. S. Swain, “Analytical distributions for stochastic gene expression,” Proceedings of the National Academy of Sciences, vol. 105, no. 45, pp. 17 256–17 261, 2008.

[55] P. Bokes and A. Singh, “Gene expression noise is affected differentially by feedback in burst frequency and burst size,” Journal of mathematical biology, vol. 74, no. 6, pp. 1483–1509, 2017.

[56] A. Singh and J. P. Hespanha, “Stochastic hybrid systems for studying biochemical processes,” Philosophical Transactions of the Royal Society A: Mathematical, Physical and Engineering Sciences, vol. 368, no. 1930, pp. 4995–5011, 2010.

[57] M. Soltani and A. Singh, “Moment analysis of linear time-varying dynamical systems with renewal transitions,” SIAM Journal on Control and Optimization, vol. 57, pp. 2660–2685, 2019.

[58] N. Friedman, L. Cai, and X. Xie, “Linking stochastic dynamics to population distribution: an analytical framework of gene expression,” Physical Review Letters, vol. 97, p. 168302, 2006.

[59] L. Cai and N. F. X. S. Xie, “Stochastic protein expression in individual cells at the single molecule level,” Nature, vol. 440, pp. 358–362, 2006.

[60] P. Bokes and A. Singh, “Controlling noisy expression through auto regulation of burst frequency and protein stability,” in International Workshop on Hybrid Systems Biology. Springer, 2019, pp. 80–97.

[61] J. P. Hespanha and A. Singh, “Stochastic models for chemically reacting systems using polynomial stochastic hybrid systems,” International Journal of Robust and Nonlinear Control, vol. 15, pp. 669–689, 2005.

[62] J. M. Pedraza and J. Paulsson, “Effects of molecular memory and bursting on fluctuations in gene expression,” Science, vol. 319, no. 5861, pp. 339–343, 2008.

[63] A. Singh and J. P. Hespanha, “Approximate moment dynamics for chemically reacting systems,” IEEE Transactions on Automatic Control, vol. 56, no. 2, pp. 414–418, 2010.

[64] I. F. Tannock, “Tumor physiology and drug resistance,” Cancer and Metastasis Reviews, vol. 20, no. 1, pp. 123–132, 2001.

[65] N. Totis, C. Nieto, A. Küper, C. Vargas-García, A. Singh, and S. Waldherr, “A population-based approach to study the effects of growth and division rates on the dynamics of cell size statistics,” IEEE Control Systems Letters, vol. 5, no. 2, pp. 725–730, 2020.

[66] C. Nieto, C. Vargas-Garcia, J. Pedraza, and A. Singh, “Cell size regulation and proliferation fluctuations in single-cell derived colonies,” bioRxiv, 20221.

[67] D. Taylor, N. Verdon, P. Lomax, R. J. Allen, and S. Titmuss, “Tracking the stochastic growth of bacterial populations in microfluidic droplets,” Physical Biology, vol. 19, no. 2, p. 026003, 2022.

